# Comparing *in vitro* cytotoxic drug sensitivity in colon and pancreatic cancer using 2D and 3D cell models: contrasting viability and growth inhibition in clinically relevant dose and repeated drug cycles

**DOI:** 10.1101/2023.11.22.568335

**Authors:** Tia R. Tidwell, Gro Røsland, Karl Johan Tronstad, Kjetil Søreide, Hanne R. Hagland

## Abstract

*In vitro* drug screening that is more translatable to the *in vivo* tumor environment can reduce both time and cost of cancer drug development. Here we address some of the shortcomings in screening and show how treatment with 5-fluorouracil (5-FU) in 2D and 3D culture models of colorectal cancer (CRC) and pancreatic ductal adenocarcinomas (PDAC) give different responses regarding growth inhibition.

The sensitivity of the cell lines at clinically relevant 5-FU concentrations was monitored over 4 days of treatment in both 2D and 3D cultures for CRC (SW948 and HCT116) and PDAC (Panc-1 and MIA-Pa-Ca-2) cell lines. The 3D cultures were maintained beyond this point to enable a second treatment cycle at day 14, following the timeline of a standard clinical 5-FU regimen. Evaluation after one cycle did not reveal significant growth inhibition in any of the CRC or PDAC 2D models. By the end of the second cycle of treatment the CRC spheroids reached 50% inhibition at clinically achievable concentrations in the 3D model, but not in the 2D model. The PDAC models were not sensitive to clinical doses even after two cycles. High content viability metrics point to even lower response in the resistant PDAC models.

This study reveals the limitations of testing drugs in 2D cancer models and short exposure in 3D models, and the importance of using appropriate growth inhibition analysis. We found that screening with longer exposure and several cycles of treatment in 3D models suggests a more reliable way to assess drug sensitivity.

## 1. Introduction

Standard cancer drug screening in its current form lacks the complexity necessary to represent key clinical aspects of human tumors. Primary *in vitro* drug screening typically takes place in two-dimensional (2D) monolayer cell cultures with growth as the primary measure of response^1,2^. However, cell culture in 2D can be quite sensitive to treatment, resulting in inflated drug response^3^ and inconsistent results^4^. Despite originally being derived from tumors, 2D culture diverges from the *in vivo* tumor environment, including lack of complex intercellular connections and saturation of nutrients. The nutrient environment changes significantly from 2D cultures to 3D culture by virtue of pure morphology and the nutrient gradients formed in 3D culture^5^. This can directly affect and be affected by cell metabolism, which has been shown to be important in drug response in both 2D^6,7^ and 3D^8,9^. Drug screening in three-dimensional (3D) spheroid cultures are more relevant and give better insight into how the cancer will respond to treatment^10^ by offering a more physiological microenvironment.

Measurement and reporting of response are other areas for improvement in drug screening. By measuring growth alone, directly or indirectly, the full effect of a drug is not assessed. A drug may be cytostatic and arrest growth of a cell line, whereby cells resume growth upon removal of the drug. Most cell stains used for drug response assays are assessing growth and reported as relative intensity values. Viability stains report directly on quantity of live cells; however, these results must also be normalized. In spheroids, analysis of response can be more challenging than the analysis of flat 2D cultures. Simple measurement of growth in 3D requires different approaches and viability staining is more complex with the addition of the third dimension. By nature, healthy spheroids may have a necrotic core leading to accumulation of stains detecting dead cells, whereas smaller spheroids may exhibit smaller necrotic areas and less dead cell stain. Incorporating subtle changes in analysis of viability staining between healthy and treated spheroids is thus important. Performing high content screening using metrics of viability can provide more information beyond standard growth inhibition measurements. For reporting growth response, it is also important to be clear if the inhibitory concentration (IC50) reported is relative IC50 or absolute IC50 (also known as GI50^11^), where relative IC50 demonstrates just 50% of the maximum effect on a particular sample and absolute IC50 is 50% inhibition compared to an untreated control^12^.

While not common as a monotherapy anymore, 5-FU is one of the oldest anti-cancer drugs and remains an integral part of standard combination therapies for CRC^13,14^ (e.g. FOLFIRI, FOLFOX) and PDAC^15,16^ (e.g. FOLFIRINOX). However, tumor responsiveness varies^17^ and adverse effects related to toxicity are a major issue^11,18^. Thus, understanding 5-FU sensitivity is important and offers a good test model for drug screening. Here we use concentrations of 5-FU that correspond to clinically relevant doses, up to 4 ug/ml^19–23^ and higher (16 ug/ml) to investigate the response of the different *in vitro* cell models. While 5-FU treatment regimens vary, it is frequently found to include multiple treatment cycles of 46-96 hour continuous infusion^20–24^. In accordance with this, we exposed 2D and 3D cultures to 5-FU for 4 days continuously, with the 3D cultures subjected to a second 4-day treatment cycle 2 weeks after the first.

The aims of this study were to compare response to common chemotherapy in cell models of colorectal cancer (CRC) and pancreatic ductal adenocarcinoma (PDAC) using an enhanced screening method and correlate to the cells’ initial metabolic phenotypes. To enable this, we treated 2D and 3D cultures with 5-fluorouracil (5-FU) over a range of clinically relevant concentrations and monitored response according to both growth and viability metrics.

## 2. Materials and Methods

### 2.1. Cell Culture

SW948 (CRC) and MIA-Pa-Ca-2 (PDAC) were purchased from European Collection of Authenticated Cell Cultures (ECACC), HCT116 (CRC) and Panc1 (PDAC) cell lines were generously provided by collaborators Laboratory for Molecular Biology at Stavanger University Hospital. All cell lines were cultured in DMEM (Corning, Corning, USA) supplemented with 10% fetal bovine serum (FBS) (BioWest, Nuaillé, France), 5 mM glucose (Sigma-Aldrich, St. Louis, USA), 2 mM L-glutamine (Corning, Corning, USA), penicillin (100 U/ml) & streptomycin (100 μg/ml) (Merck Millipore Corporation, Burlington, USA) in a humidified incubator at 37°C with 5% CO2 infusion. Cells were grown in 2D adherent culture conditions, of which spheroids were prepared before each experiment. For 2D experiments, cells were seeded in flat-bottom 96-well plate (VWR, Radnor, USA) at a density of 5 000 cells/well. Cells attached overnight and were treated the following day. Spheroids were produced from 40 μl volumes of detached single-cell suspensions at 1.25x10^5^ cells/ml in CELLSTAR cell repellent U-bottom plates (Greiner Bio-One, Kremsmünster, Austria). Spheroids were grown for 3 days (CRC) or 4 days (PDAC) before 5-fluorouracil treatment.

### 2.2. Drug treatment

5-fluorouracil powder (EMD Millipore Corp., Bellerica, USA) was reconstituted in water at a concentration of 1 mg/ml. Further dilutions to achieve the desired treatment concentrations were made in growth media and 100 ul was added to each well. The cells were exposed to the treatment for 4 days. In 2D, this marked the end of the experiment. For spheroids, the media was exchanged on day 4 and every other day thereafter, until day 14 (from initial drug exposure), when a new treatment was applied. On day 18, the media was again exchanged to remove the treatment and they were maintained for 3 additional days for post-treatment imaging.

### 2.3. Imaging and image analysis

All samples were imaged using a Leica SP8 confocal microscope and 5X and 20X dry objectives, for spheroids and 2D cultures, respectively. Regular monitoring of growth in 2D and 3D occurred every day by capturing transmitted light images during treatment rounds, immediately preceding treatment on days 7 and 11 between the two rounds, and day 21 after the second round. Confluency of growth in 2D cultures was measured in the images using the PHANTAST^25^ MATLAB interface. Spheroid area was measured in the images using ImageJ^26^/FIJI^27^ macros: SpheroidArea (Additional File 1) and SpheroidArea2 (Additional File 2). Calcein AM (Biotium, Fremont, USA; final concentration in well: 4 μg/ml, to stain living cells) and propidium iodide (Sigma-Aldrich, St. Louis, USA; final concentration in well: 7.5 μg /ml, to stain dead cells) was added to 5 wells from each cell line on days 3 and 17 (before the end of each round) to incubate overnight for imaging the following day. Z-stacks of fluorescent images were captured (CAM, Ex: 488 nm, Em: 493-529 nm; PI, Ex: 552 nm, Em: 630-643 nm). Images were processed and analyzed using ImageJ/FIJI macros: StackProfleData2 (Additional File 3) and StackProfileData3 (Additional File 4). In short, plot profiles of a selected rectangular area that spanned the spheroid center were taken for each image and channel in the stack (Figure S2, Additional File 5).

### 2.4. Data analysis

#### 2.4.1. Spheroid Growth

Spheroid area measurements (mm^2^) were combined in R and analyzed in GraphPad Prism (Version 9). Spheroid diameter (d, microns) was estimated from area (A) using the following equation: d=2000· √(A/π). The sizes of the treated spheroids at each timepoint were baselined to the size of the control on the same day, yielding percent of control (charts containing raw values can be found in Figure S1, Additional File 5). A final value percentage was constructed by an average across experiments. 2-way ANOVA was run to establish statistical significance (Tukey test for multiple comparisons). Percent growth inhibition over the 5-FU treatment range was used for nonlinear regression analysis and calculation of GI50 using GraphPad’s Absolute IC50 equation.

#### 2.4.2. Viability staining

Plot profiles of viability staining were exported from ImageJ and imported into GraphPad Prism (Version 9). Stack profiles were added together and a rolling average of 3 was taken to reduce noise in peak analysis. The area under the curve (AUC) was taken (baseline at lower quartile) and this data was exported for consolidation and analysis in R. Metrics were calculated and extracted from the AUC data including the total AUC, baseline for each channel, maximum peak heights, width of widest peaks, and distances between peaks, resulting in a total of 14 variables (Figure S1, Additional File 5).

#### 2.4.3. Principal Component Analysis

Changes in parameters over concentration varied between the cell lines and days in such a way that a pattern was not easily discernible, so principal component analysis (PCA) was used to extract response from all variables together. PCA was performed in GraphPad Prism (Version 9) using 14 variables and selection of the two principal components with the largest eigenvalues (Figure S3, Additional File 5). Data was scaled to have a mean of 0 and SD of 1. After PCA, any variable with loadings less than 0.5 were excluded to preserve the highest contributing metrics. The final PCA included 9 variables. Mean values for each principal component (PC) were calculated and the Euclidean distance (d) between each mean point of a treatment from the mean control position was calculated using the following equation:

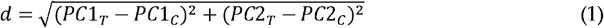

Response was grouped according to mean distance from control: low response is a distance of 0.432 or below (25^th^ percentile), intermediate response between 0.433 and 1.364 (interquartile range), and high response equal to or greater than 1.365 (75^th^ percentile).

## 3. Results

### 3.1. Longer 5-FU exposure is required in 3D cultures to reach GI50 compared to 2D cultures

The sensitivity of the cell lines at clinically relevant 5-FU concentrations was monitored over 4 days of treatment in both 2D and 3D cultures (Figure 1A). The 3D cultures were maintained beyond this point to enable a second treatment cycle at day 14, following the timeline of a standard clinical 5-FU regimen. At clinically achievable and safer concentrations of 5-FU (up to 4 μg/ml), only HCT116 growth is inhibited beyond 50% upon 4 days of treatment exposure (Figure 1B). This level of inhibition is not achieved in 3D culture of HCT116 at day 4. Higher concentration than 4 μg/ml of 5-FU does not increase inhibition dose-dependently, implying a maximum effect has been reached. While the other cell lines in 2D do not reach 50% inhibition by day 4, they demonstrate significant inhibition compared to untreated controls and lower concentrations of 5-FU (0.25-0.5 μg/ml) and exhibit more inhibition than their corresponding 3D cultures at the same timepoint. By day 18, HCT116 is still the most sensitive with spheroids treated with 1 μg/ml, reaching nearly 70% inhibition of growth. SW948 also reaches 50% growth inhibition by day 18 in 1 μg/ml and higher concentrations. Minimal additional inhibition is also seen in cells treated with > 4 μg/ml in the SW948 spheroids at day 18. In PDAC spheroids, 50% inhibition is only reached at 16 μg/ml, far beyond a safe clinical concentration. Notably, in MIA-Pa-Ca-2, growth resumes after removal of 5-FU in these lower concentrations.

**Figure 1.**
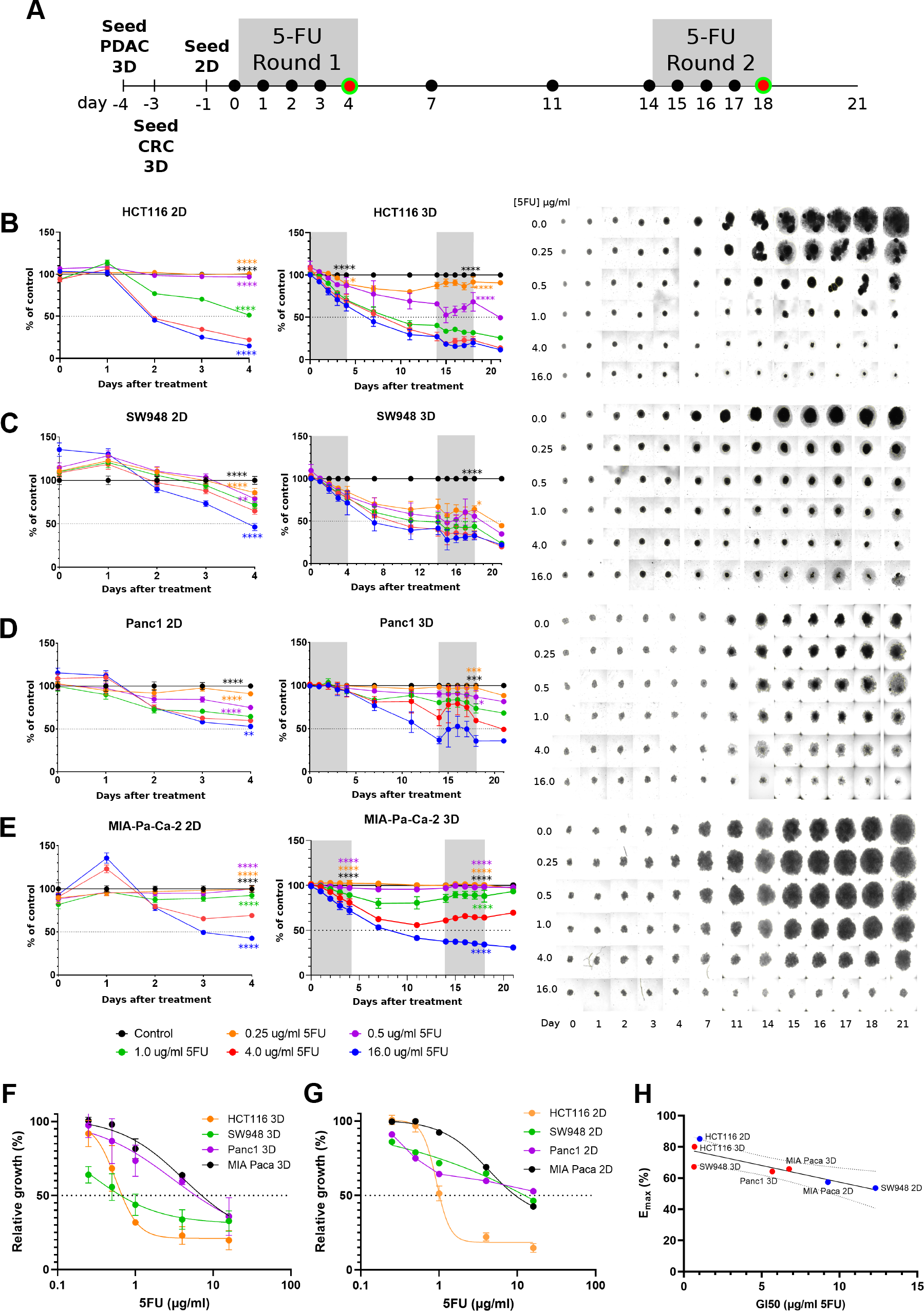
Treatment and response of 2D and 3D cultures with 5-FU. (A) Timeline of 5-FU treatment, where black circles indicate brightfield images taken to monitor growth and red/green circles indicate viability imaging. Growth results over the treatment period, from right to left showing 2D results, 3D results, and 3D images for each cell line: (A) HCT116, (B) SW948, (C) PANC1, (D) MIA-Paca-2. Error bars represent SEM. N=40 over 2 independent experiments. 2-way ANOVA of 4 μg/ml results versus other concentrations at day 4 (2D and 3D) and day 18 (3D): *, p<0.05. Dose-response curves of (F) 3D at day 18, and (G) 2D cultures at day 4, used for calculation of GI50. Error bars represent SD. (H) Correlation of GI50 with Emax: -0.8648, p=0.012, R^2^ = 0.7475. Panc1 2D was not inhibited to 50%.

Growth relative to untreated control samples was plotted against concentration of 5-FU to estimate the maximum effect (Emax) and concentration of 5-FU to reach 50% growth inhibition (GI50) (Figure 1F-G). This provides a quantification of the relative sensitivities of the cultures over the treatment period and more specific insight into what concentration is needed to reach 50% inhibition. GI50 of HCT116 in 2D culture by day 4 is 1.01 mg/ml and for 3D culture on day 18 is 0.67 mg/ml. For SW948, GI50 for 2D and 3D cultures are 12.3 mg/ml and 0.65 mg/ml, respectively. Lower response is seen in the PDAC cell lines, and this is reflected in the GI50 values. Panc1 GI50 for 3D culture is 5.68 mg/ml, while 2D culture did not reach 50% inhibition by day 4 so GI50 is not possible to calculate. MIA-Pa-Ca did reach 50% inhibition in 2D, yielding a GI50 of 9.25 mg/ml; GI50 of 3D culture is 6.76 mg/ml. Emax and GI50 correlation is -0.8648, p=0.0120 (Figure 1H). None of the cell lines reached 50% inhibition by day 4 in 3D culture.

### 3.2. High content analysis of drug response in 3D cultures requires combined viability metrics

Spheroids were stained with calcein acetoxymethyl (calcein AM or CAM) and propidium iodide (PI) to assess location and proportion of viable and dead cells, respectively, and imaged on days 4 and 18 (at the end of each treatment round) (Figure 2). Live/dead staining in spheroids is more complex than reporting the relative values of the stain intensity. As can be seen just from simple visual assessment, even control spheroids have a large amount of PI staining indicating a necrotic core, and the intensity is high due to the overall larger size of the spheroid. It is also apparent that the viable cells persist to some degree in all the cultures over the treatment period. Using the plot profiles of the two stains and transmitted light, the area under the curve (AUC) of each was analyzed and used to produce several different viability metrics (Figure S2, Additional File 5). The stains followed different patterns depending on the cell line; a simplified view of this, using untreated controls as an example, is visualized in Figure 4. Viable cells remain in all cell lines, even at the end of treatment, but the percent of dead cells (PI core) or light-impermeable core (TL core) increases in all cell lines.

**Figure 2.**
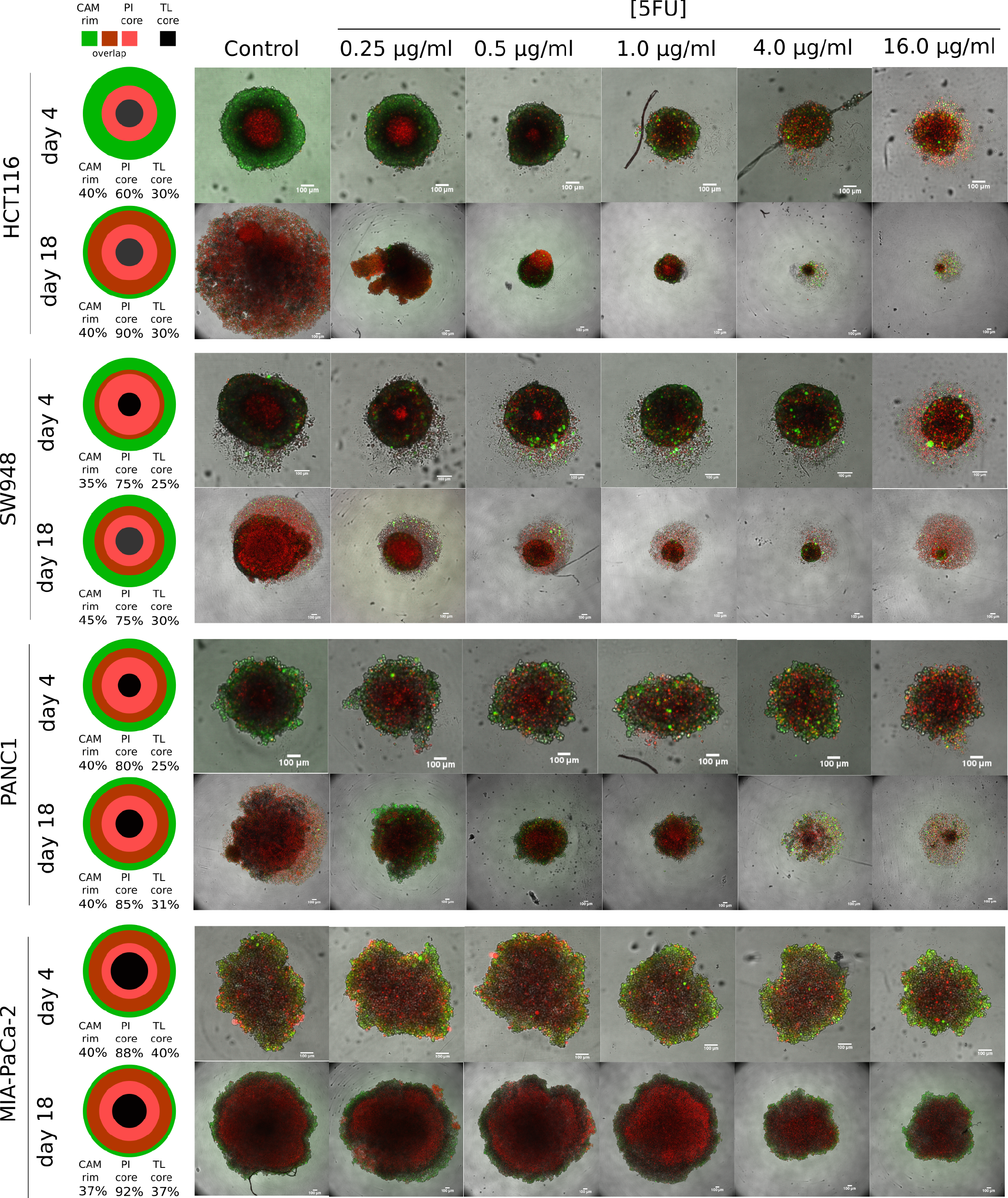
Viability staining of spheroids. In the leftmost column, measurements of CAM rim, PI core and TL core in control spheroids of each cell line are given as the average percent of spheroid diameter (Figure S2), at each viability imaging timepoint. The size of illustration is not indicative of spheroid size, but to represent relative proportions of staining in each so they are all equal. To the right, representative composite images of viability staining of spheroids on days 4 and 18 of 5-FU treatment (actual n=5). Images are maximum projections of a stack of images. Dead cell staining: propidium iodide (red), Viable cell staining: calcein AM (green). 100 μm scale bar in white. Abbreviations: calcein AM, CAM; propidium iodide, PI; transmitted light, TL.

As these and other parameters vary between cell lines, a clear pattern was difficult to distinguish. Using principal component analysis, the parameters were combined and those that differentiated them the most were grouped together into principal components (Figure S3-S4, Additional File 5). In PC1 (capturing 39.6% of variance), the AUC per micron of all stains (loading from 0.648-0.801), CAM peak intensity (0.843), PI peak intensity (0.799), and CAM core (−0.631) are the heaviest weighted values. In PC2, the defining values are the staining baseline values (CAM: 0.714, PI: 0.734, TL: 0.632). Plotting the scores of each individual sample from each component reveals some grouping patterns (Figure 3). The day 4 and day 18 samples cluster differentially (Figure 3A), except MIA-Pa-Ca-2 at day 18 also appears within the day 4 region. As most of the day 18 values are clustered in positive PC1 region, this points to higher AUC per micron and higher PI peak, among other smaller parameter contributions. Lower 5-FU concentrations are characterized by positive PC2 values, which is heavily influenced by either a large core % (as measured by CAM), low baseline values, or a combination of both. Figure 3B shows how the cell lines differ using these parameters (See Figure S5 in Additional File 5 for separate charts of each cell line). Higher concentrations of 5-FU and day 18 values appear a greater distance from the day 4 and lower concentration values in CRC cell lines. Overall, the cell lines follow different spread patterns of PC scores.

**Figure 3.**
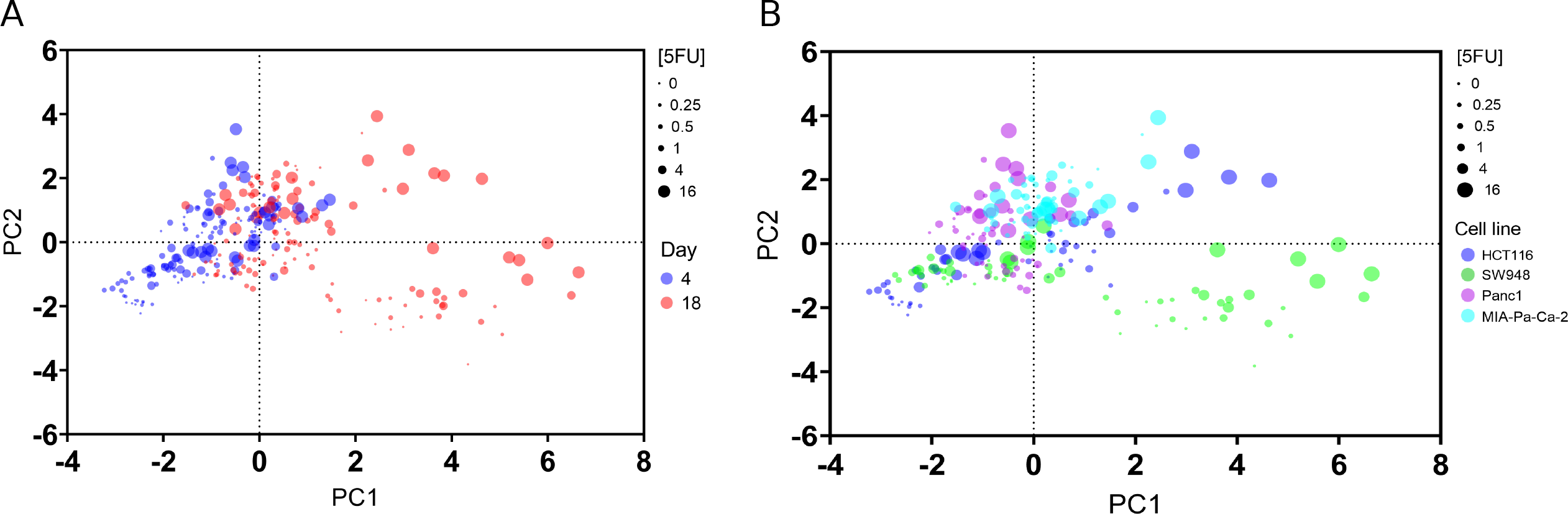
Samples clustered by principal component scores from viability data. (A) Samples plotted by principal component scores, colored by **day of analysis**. Size of dots vary by concentration of 5-FU, from lowest = smallest dot to highest concentration = largest dot. (B) Samples plotted by principal component scores, colored **by cancer cell line**. Size of dots vary by concentration of 5-FU, from lowest = smallest dot to highest concentration = largest dot.

The distance of treatments from the control group of each cell line (Table 1) allows quantification of response between the different cell models and endpoints. HCT116 is the most responsive to treatment with the largest change using this viability measurement as well, and in order of decreasing response are SW948, Panc1, and MIA-Pa-Ca-2. This mirrors the growth measurements but amplifies the lack of change in the low-response MIA-Pa-Ca-2, where very little change is seen in viability metrics.

**Table 1.** Principal component score mean distances from untreated control group, by concentration, as an indication of response to 5-FU. Bold and underline: high response, greater than 1.365. Bold only: intermediate response, between 0.433 and 1.364. Italics: low response, less than or equal to 0.432.

| [5-FU] (µg/ml) | Day 4 (post-treatment 1°) |  |  |  |  | Day 18 (post-treatment 2°) |  |  |  |  |
| --- | --- | --- | --- | --- | --- | --- | --- | --- | --- | --- |
|  | 0.25 | 0.5 | 1.0 | 4.0 | 16.0 | 0.25 | 0.5 | 1.0 | 4.0 | 16.0 |
| HCT116 | *0.122* | **0.725** | **1.007** | **<u>1.464</u>** | **<u>2.137</u>** | **0.724** | **0.747** | **0.598** | **<u>1.465</u>** | **<u>4.222</u>** |
| SW948 | *0.312* | **0.903** | **0.592** | *0.265* | **<u>1.539</u>** | **0.812** | **<u>1.560</u>** | **1.317** | **<u>1.516</u>** | **<u>3.440</u>** |
| Panc1 | *0.401* | **0.630** | **0.433** | **0.863** | **<u>1.957</u>** | *0.430* | **0.630** | **0.574** | **<u>1.880</u>** | **1.332** |
| MIA-Pa-Ca-2 | *0.263* | *0.191* | *0.419* | *0.415* | **1.065** | **0.787** | **1.056** | **0.705** | **1.134** | *0.268* |

### 3.3. Effect of cellular metabolism

The metabolic phenotype of these cell lines in 2D and 3D cultures have been established previously^28^. To assess whether the metabolic phenotype, presented as the ratio of oxygen consumption rate (OCR) to extra-cellular acidification rate (ECAR), influenced the drug response the correlation of 5-FU Emax to OCR/ECAR was calculated. Although not statistically significant, we found that Emax is negatively correlated with OCR/ECAR (−0.689, p=0.059) (Figure 4) across 2D and 3D growth settings.

**Figure 4.**
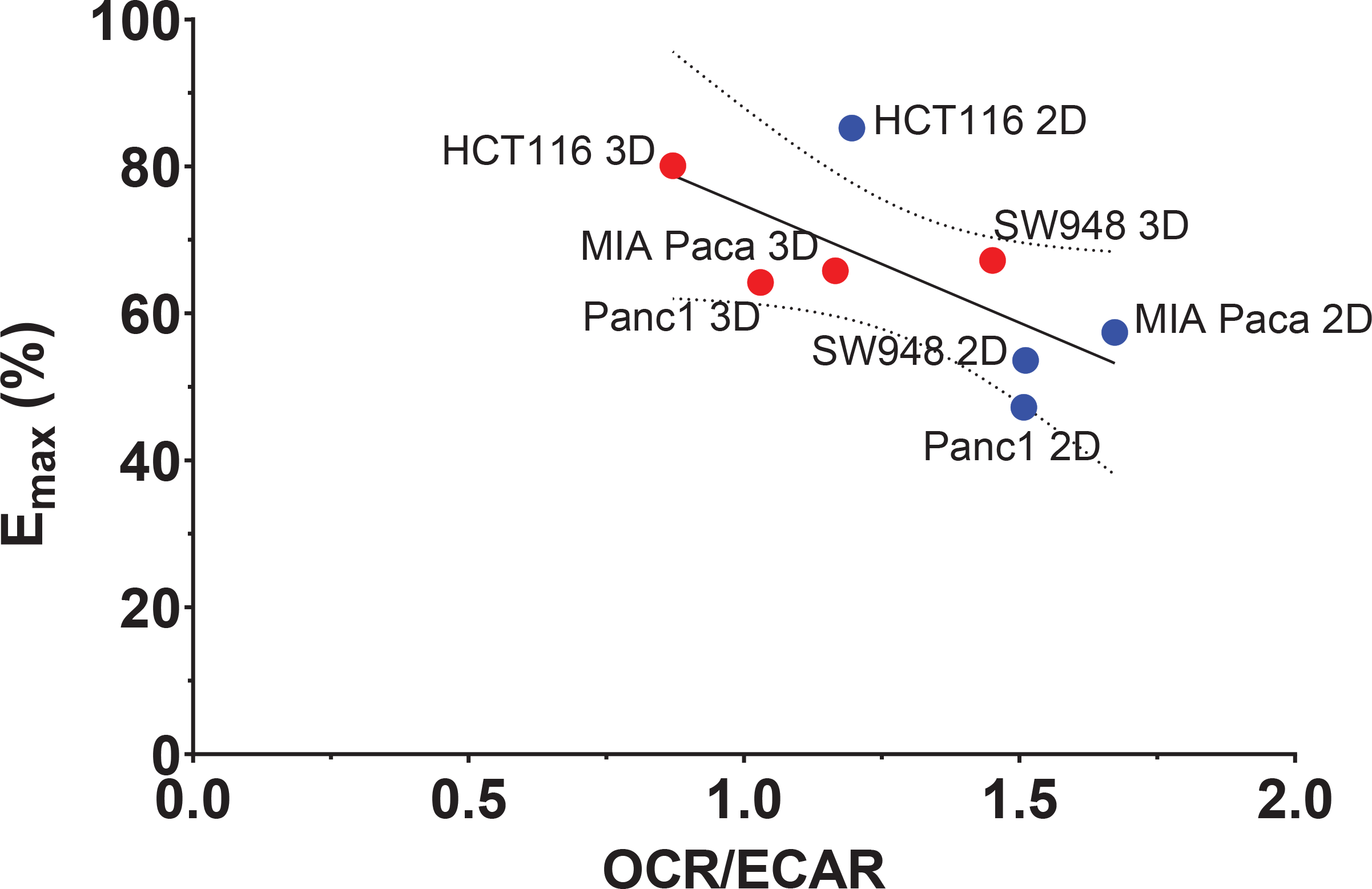
Correlation of Emax to metabolic phenotype indicator, OCR/ECAR. Emax is negatively correlated with OCR/ECAR (−0.689, p=0.059, R^2^ = 0.4752). OCR: oxygen consumption rate, ECAR: extracellular acidification rate. OCR/ECAR data (Table S1) from Tidwell *et al*^28^.

## 4. Discussion

The importance of testing drug sensitivity of cancer models in 3D cultures is emphasized in this study. 2D cultures exhibit increased sensitivity to 5FU treatment when compared to 3D cultures, both in terms of maximal growth inhibition and concentration required to achieve 50% growth inhibition. Typical 2D culture also lacks the possibility for longer culture timelines to understand multi-round drug response and resistance. We found that the relative effects of the 5-FU concentrations in 2D are somewhat predictive of the relative effects in 3D, which could give an idea of the most active range of 5-FU to apply in 3D. However, the 5-FU response in 3D is a comparison of results at day 18 to day 4 of 5-FU 2D response metrics. This could be conceived as misleading, however at day 4 treatment effect in 3D were minimal and did not reach 50% inhibition in any of the cell lines tested. It is clear from these results that short-term studies with spheroids have limited quantitative utility when using relevant dosing. The increased sensitivity of 2D cultures over 3D cultures was expected in line with other studies showing 3D drug response^29,30^. Each cell in 2D is being exposed to much more drug in culture, with the surface area of cells in a monolayer much larger than that in spheroids. This is due to both the organization and morphology of the cells in the 3D structure, with many cells located in the interior of the spheroid. The spheroid also experiences diffusive exposure to the drug beyond the surface, due to naturally occurring gradients^31^.

The response metrics used from growth measurements here, Emax and GI50, have some disadvantages^32^. Despite being widely used, these metrics may be highly correlated to cell doubling time and other environmental conditions^32^. Here, the GI50 numbers are calculated using relative growth to respective controls, which should reduce the effect of cell doubling times between cell lines. Consequently, the doubling times of these cell lines^33^ are not correlated with the GI50 values, supported by our findings. HCT116 is the fastest growing cell line (17.9 hours) and does have the lowest GI50, but the other cell lines have similar 2D doubling times (between 24-28 hours) and vary considerably in 2D and 3D GI50 values (doubling times in 3D were not calculated or considered). Additionally, it could be argued that sensitivity to a drug based on cell division rate and other culture conditions can be important variables in drug response to consider and not ignore. Regardless, the general limitations of these metrics are known, so additional work with viability staining here offers expanded insight and an approach for high content screening. While there may be significant inhibition of growth between concentrations, viability may not differentiate them as much in some cases, as seen in MIA-Pa-Ca-2. The staining also reveals the challenge of using viability stains in spheroids. For example, there is a high amount of PI staining in the untreated spheroids due to cell death in the core. Therefore, using a measurement of staining intensity alone can be misleading without considering the size of the spheroid. Furthermore, there are regions that are co-stained with both calcein AM and PI, possibly due internalization of dyes in the cells, or it could be caused by a mix of cells in these regions which are overlapping. Since calcein AM was stained overnight, cells that were viable at the start could have progressed to cell death and membrane disruption causing additional PI staining to occur. This is a challenge when working with 3D spheroids, requiring longer incubation when using dyes to achieve sufficient diffusion into the spheroid, whereby resolution of cell death may be incorrectly interpreted. Nonetheless, these studies are important to acquire knowledge on the structure of spheroids using different staining which can be useful in further modeling spheroid growth and drug response *in silico*. The results from the viability metrics were able to differentiate response in the spheroid models without size serving as a distinct variable. However, size was incorporated and corrected for by having AUC/micron and core size as a percent of the spheroid diameter. Other variables such as baseline and peak values were completely size independent.

5-FU is commonly used in combination with other drugs to enhance cancer treatment response^34–36^. There is a high variability in reported measured plasma concentrations of this drug, whereby most studies have found average steady state levels to be about 1-6 μg/ml, but values range from 0.3 to 60 μg/ml^19–23^. Overall, the plasma levels are linearly correlated to g/m^2^ dosing^21^, where an increase of 500 mg/m^2^ 5-FU, may cause an increase in the plasma concentration of 200 ng/ml. The recommended dose is 2.5-3 μg/ml for optimal balance of tumor response and systemic toxicity^21^, however this was found not be an effective treatment concentration in the PDAC spheroids. A possible explanation to drug insensitivity is how the metabolism of cancer cells are primed to either glycolysis or OXPHOS, as we have found previously^33^. This has been well documented in 2D cancer grown cancer cells^6,7^, but also in 3D spheroids^8,9^. A common measure of metabolic phenotype is the OCR/ECAR ratio, OCR or oxygen consumption is a measure of glucose metabolism via oxidative phosphorylation in the mitochondria and extracellular acidification is proportional to lactate excreted from glycolytic metabolism of glucose. Therefore, a higher OCR/ECAR ratio denotes a more oxidative metabolic phenotype and a lower ratio, a more glycolytic phenotype. The correlation here of Emax with OCR/ECAR is not statistically significant but is an interesting finding, whereby a high OCR/ECAR ratio or more oxidative phenotype correlates with lower maximum inhibition by 5-FU. Expansion of the screening to more cell lines is essential to validate this correlation. Perhaps in contrast, *Zhao et al*. have shown 5-FU resistant cells increase glucose uptake and glycolytic activity^37^. While we find here that a more glycolytic phenotype correlates to higher sensitivity, this is the pre-treatment phenotype. The study by Zhao *et al*.^37^ presents metabolic changes from treatment, which is something that should be done with these cell lines as well. In the models that are more resistance to treatment, pre-treatment with a metabolic drug or combination treatment should be investigated. In previous work, an oxidative CRC cell line (in 2D), metformin has been shown to shift metabolism to a more glycolytic phenotype^6^. Whether this is reproduced in 3D culture, and extends to other oxidative cell lines, needs to be investigated. Conversely, dichloroacetate (DCA), an inhibitor of pyruvate dehydrogenase kinase (PDK1), has been shown to be a possible candidate to shift metabolism away from glycolysis^37,38^.

The culture model and treatment approach here is simplified in order to study the longer screening time-line and isolate response. Other drugs used in current clinical care as combination therapies with 5-FU such as folinic acid, irinotecan, oxaliplatin, and gemcitabine should be included in future experiments, particularly for PDAC studies.

## 5. Conclusions

We find here that not only is 3D culture essential for assessing drug sensitivity in these *in vitro* cancer models, compared to 2D, but also longer multi-cycle treatments are achievable and further informative of response. Using only four cell models, we can distinguish between models that exhibit high and low 5-FU sensitivity. Both CRC cancer models are more sensitive to 5-FU than the PDAC models, as expected from clinical experience. We present a model that can be used for variety of applications and provides a low barrier of entry at the simplest level of 3D culture techniques. The highest impact of application would come from culturing primary cells and treating them, knowing the matched patient outcomes from actual treatment, leading to robust personalized medicine and improved screening for drug discovery.

## Supporting information

StackProfileData3

StackProfileData2

SpheroidArea

SpheroidArea2

Viability metrics

## Author Contributions

Conceptualization, T.T. and H.R.H, investigation and experimental set up T.T., writing—original draft preparation, T.T., H.R.H., writing—review and editing, all authors. All authors have read and agreed to the published version of the manuscript.

## Data Availability Statement

The datasets generated during the current study are available on figshare, https://doi.org/10.6084/m9.figshare.16702204.v1

## Acknowledgments

We would like to thank head lab engineer Julie Nikolaisen (PhD) for lab assistance and Stavanger University Hospital for providing us with the cell lines used in these experiments.

## Supplementary Materials

- Additional File 1: ImageJ macro SpheroidArea (text file)
- Additional File 2: ImageJ macro SpheroidArea2 (text file)
- Additional File 3: ImageJ macro StackProfileData2 (text file)
- Additional File 4: ImageJ macro StackProfileData3 (text file)
- Additional File 5: Supplementary Figures S1-S5, Table S1 (Word file)
- Additional File 6: Raw values of viability metrics used for principal component analysis. Cell lines are numbered as follows: HCT116, 1; SW948, 2; Panc-1, 3; MIA-Pa-Ca-2, 4. (csv file)

## References

1. NCI-60 Human Tumor Cell Lines Screen | Discovery & Development Services | Developmental Therapeutics Program (DTP). Available at https://dtp.cancer.gov/discovery_development/nci-60/default.htm (2021).

2. Shoemaker, R. H. The NCI60 human tumour cell line anticancer drug screen. Nature reviews. Cancer 6, 813–823; 10.1038/nrc1951 (2006).

3. Tung, Y.-C. et al. High-throughput 3D spheroid culture and drug testing using a 384 hanging drop array. The Analyst 136, 473–478; 10.1039/c0an00609b (2011).

4. Haibe-Kains, B. et al. Inconsistency in large pharmacogenomic studies. Nature 504, 389–393; 10.1038/nature12831 (2013).

5. Sutherland, R. M. et al. Oxygenation and differentiation in multicellular spheroids of human colon carcinoma. Cancer research 46, 5320–5329 (1986).

6. Alhourani, A. H. et al. Metformin treatment response is dependent on glucose growth conditions and metabolic phenotype in colorectal cancer cells. Sci Rep 11; 10.1038/s41598-021-89861-6 (2021).

7. Gui, D. Y. et al. Environment Dictates Dependence on Mitochondrial Complex I for NAD+ and Aspartate Production and Determines Cancer Cell Sensitivity to Metformin. Cell metabolism 24, 716–727; 10.1016/j.cmet.2016.09.006 (2016).

8. Russell, S., Wojtkowiak, J., Neilson, A. & Gillies, R. J. Metabolic Profiling of healthy and cancerous tissues in 2D and 3D. Scientific reports 7, 15285; 10.1038/s41598-017-15325-5 (2017).

9. Fan, T. W.-M. et al. Stable Isotope-Resolved Metabolomics Shows Metabolic Resistance to Anti-Cancer Selenite in 3D Spheroids versus 2D Cell Cultures. Metabolites 8; 10.3390/metabo8030040 (2018).

10. Gilazieva, Z., Ponomarev, A., Rutland, C., Rizvanov, A. & Solovyeva, V. Promising Applications of Tumor Spheroids and Organoids for Personalized Medicine. Cancers 12; 10.3390/cancers12102727 (2020).

11. Begg, S. K. S. et al. FOLFIRINOX Versus Gemcitabine-based Therapy for Pancreatic Ductal Adenocarcinoma: Lessons from Patient-derived Cell Lines. Anticancer research 40, 3659–3667; 10.21873/anticanres.14355 (2020).

12. Sebaugh, J. L. Guidelines for accurate EC50/IC50 estimation. Pharmaceutical statistics 10, 128–134; 10.1002/pst.426 (2011).

13. Dekker, E., Tanis, P. J., Vleugels, J. L. A., Kasi, P. M. & Wallace, M. B. Colorectal cancer. The Lancet 394, 1467–1480; 10.1016/S0140-6736(19)32319-0 (2019).

14. Brouwer, N. P. M. et al. An overview of 25 years of incidence, treatment and outcome of colorectal cancer patients. International journal of cancer 143, 2758–2766; 10.1002/ijc.31785 (2018).

15. Mizrahi, J. D., Surana, R., Valle, J. W. & Shroff, R. T. Pancreatic cancer. The Lancet 395, 2008–2020; 10.1016/S0140-6736(20)30974-0 (2020).

16. McGuigan, A. et al. Pancreatic cancer: A review of clinical diagnosis, epidemiology, treatment and outcomes. World journal of gastroenterology 24, 4846–4861; 10.3748/wjg.v24.i43.4846 (2018).

17. Mader, R. M., Müller, M. & Steger, G. G. Resistance to 5-Fluorouracil. General Pharmacology: The Vascular System 31, 661–666; 10.1016/S0306-3623(98)00191-8 (1998).

18. Tomiak, A. et al. Standard dose (Mayo regimen) 5-fluorouracil and low dose folinic acid: prohibitive toxicity? American journal of clinical oncology 23, 94–98; 10.1097/00000421-200002000-00025 (2000).

19. Terret, C. et al. Dose and time dependencies of 5-fluorouracil pharmacokinetics. Clinical pharmacology and therapeutics 68, 270–279; 10.1067/mcp.2000.109352 (2000).

20. Casale, F. et al. Plasma concentrations of 5-fluorouracil and its metabolites in colon cancer patients. Pharmacological research 50, 173–179; 10.1016/j.phrs.2004.01.006 (2004).

21. Blaschke, M. et al. Measurements of 5-FU Plasma Concentrations in Patients with Gastrointestinal Cancer: 5-FU Levels Reflect the 5-FU Dose Applied. JCT 03, 28–36; 10.4236/jct.2012.31004 (2012).

22. Etienne, M. C. et al. Co-variables influencing 5-fluorouracil clearance during continuous venous infusion. A NONMEM analysis. European journal of cancer (Oxford, England : 1990) 34, 92–97; 10.1016/s0959-8049(97)00345-6 (1998).

23. Fety, R. et al. Clinical impact of pharmacokinetically-guided dose adaptation of 5-fluorouracil: results from a multicentric randomized trial in patients with locally advanced head and neck carcinomas. Clinical cancer research : an official journal of the American Association for Cancer Research 4, 2039–2045 (1998).

24. Matsumoto, H. et al. Fluctuation in Plasma 5-Fluorouracil Concentration During Continuous 5-Fluorouracil Infusion for Colorectal Cancer. Anticancer research 35, 6193–6199 (2015).

25. Jaccard, N. et al. Automated method for the rapid and precise estimation of adherent cell culture characteristics from phase contrast microscopy images. Biotechnology and bioengineering 111, 504–517; 10.1002/bit.25115 (2014).

26. Caroline A. Schneider, Wayne S. Rasband & Kevin W. Eliceiri. NIH Image to ImageJ: 25 years of Image Analysis.

27. Schindelin, J. et al. Fiji: an open-source platform for biological-image analysis. Nature methods 9, 676–682; 10.1038/nmeth.2019 (2012).

28. Tidwell, T. R., Røsland, G. V., Tronstad, K. J., Søreide, K. & Hagland, H. R. Metabolic flux analysis of 3D spheroids reveals significant differences in glucose metabolism from matched 2D cultures of colorectal cancer and pancreatic ductal adenocarcinoma cell lines. Cancer & metabolism 10, 9; 10.1186/s40170-022-00285-w (2022).

29. Folkesson, E. et al. High-throughput screening reveals higher synergistic effect of MEK inhibitor combinations in colon cancer spheroids. Scientific reports 10, 11574; 10.1038/s41598-020-68441-0 (2020).

30. Mathews Griner, L. A. et al. Large-scale pharmacological profiling of 3D tumor models of cancer cells. Cell death & disease 7, e2492; 10.1038/cddis.2016.360 (2016).

31. Kasinskas, R. W., Venkatasubramanian, R. & Forbes, N. S. Rapid uptake of glucose and lactate, and not hypoxia, induces apoptosis in three-dimensional tumor tissue culture. Integrative biology : quantitative biosciences from nano to macro 6, 399–410; 10.1039/c4ib00001c (2014).

32. Hafner, M., Niepel, M., Chung, M. & Sorger, P. K. Growth rate inhibition metrics correct for confounders in measuring sensitivity to cancer drugs. Nature methods 13, 521–527; 10.1038/nmeth.3853 (2016).

33. Tidwell, T., Røsland, G. V., Tronstad, K. J., Søreide, K. & Hagland, H. R. Metabolic flux analysis of 3D spheroids reveals significant differences in glucose metabolism from matched 2D cultures of colorectal cancer and pancreatic ductal adenocarcinoma cell lines. (accepted). BMC Cancer & Metabolism; 10.1186/s40170-022-00285-w (2022).

34. Wolfe, A. R. et al. Neoadjuvant-modified FOLFIRINOX vs nab-paclitaxel plus gemcitabine for borderline resectable or locally advanced pancreatic cancer patients who achieved surgical resection. Cancer medicine 9, 4711–4723; 10.1002/cam4.3075 (2020).

35. Xie, P. et al. Pharmacogenomics of 5-fluorouracil in colorectal cancer: review and update. Cellular oncology (Dordrecht) 43, 989–1001; 10.1007/s13402-020-00529-1 (2020).

36. Conroy, T. et al. FOLFIRINOX or Gemcitabine as Adjuvant Therapy for Pancreatic Cancer. The New England journal of medicine 379, 2395–2406; 10.1056/NEJMoa1809775 (2018).

37. Zhao, J.-G., Ren, K.-M. & Tang, J. Overcoming 5-Fu resistance in human non-small cell lung cancer cells by the combination of 5-Fu and cisplatin through the inhibition of glucose metabolism. Tumour biology : the journal of the International Society for Oncodevelopmental Biology and Medicine 35, 12305–12315; 10.1007/s13277-014-2543-3 (2014).

38. Liang, Y. et al. Dichloroacetate restores colorectal cancer chemosensitivity through the p53/miR-149-3p/PDK2-mediated glucose metabolic pathway. Oncogene 39, 469–485; 10.1038/s41388-019-1035-8 (2020).

