## Supplementary Figures S1-S5, Table S1 for "Comparing *in vitro* cytotoxic drug sensitivity in colon and pancreatic cancer using 2D and 3D cell models: contrasting viability and growth inhibition in clinically relevant dose and repeated drug cycles"

**Supplementary figures and tables**

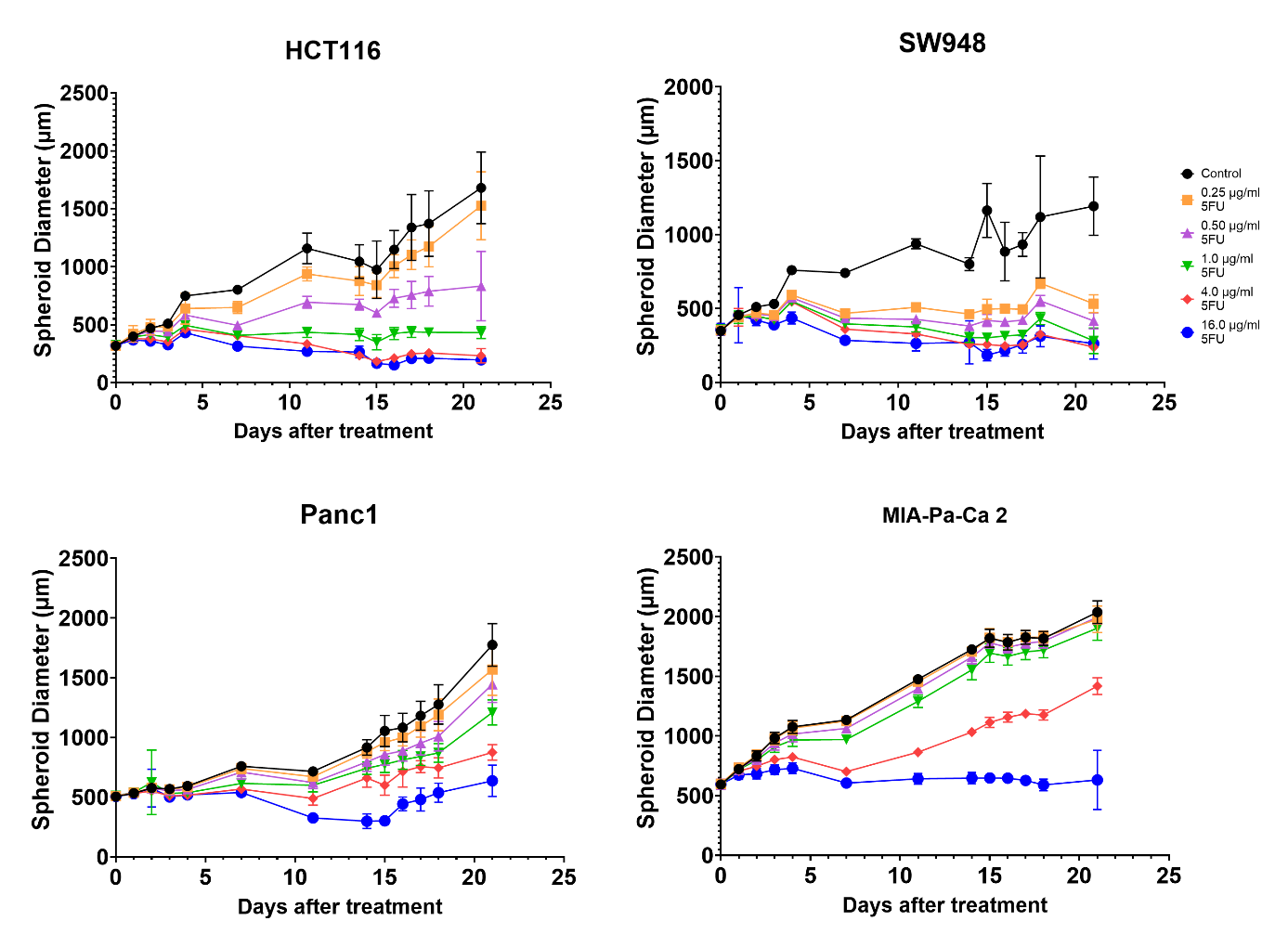

**Figure S1.** Raw size of spheroids

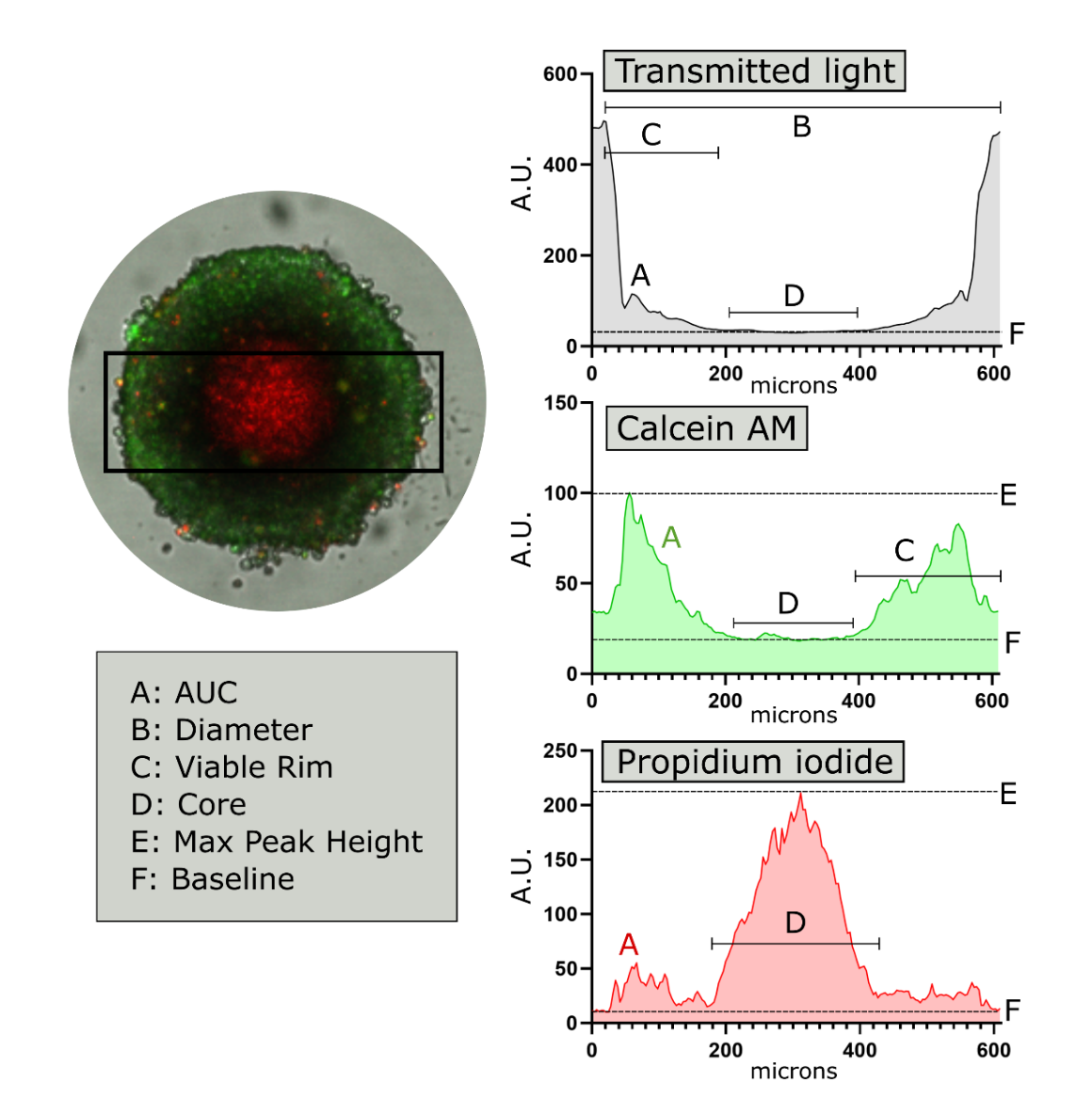

**Figure S2.** Details of metrics extracted from transmitted light (TL), calcein AM (CAM), and propidium iodide (PI) plot profiles. Example region of interest for profiles shown as black rectangle intersecting the spheroid center. A, Area under the curve (AUC). B, spheroid diameter measured from maximum distance between peaks in TL profile. C, Viable rim measured from maximum peak width in TL and CAM profiles. D, spheroid core measured from maximum distance between peak bases for TL and CAM core measurement and maximum peak width for PI core measurement. E, Max peak height measured as maximum peak height for CAM and PI profiles. F, baseline used in AUC analysis which is at the lower quartile or 25% of the profile data amplitude. Only peaks beyond 10% over the baseline were included in analysis. Abbreviations: area under the curve, AUC; arbitrary units, A.U.

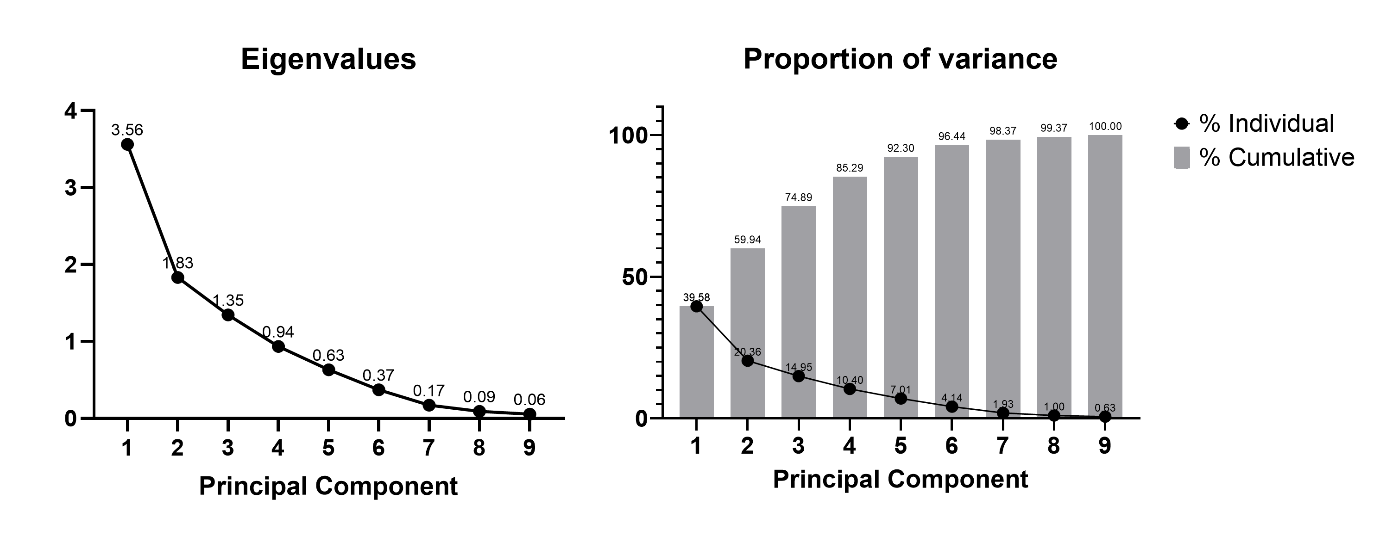

**Figure S3.** Eigenvalues and proportion of variance for principal components in the PCA.

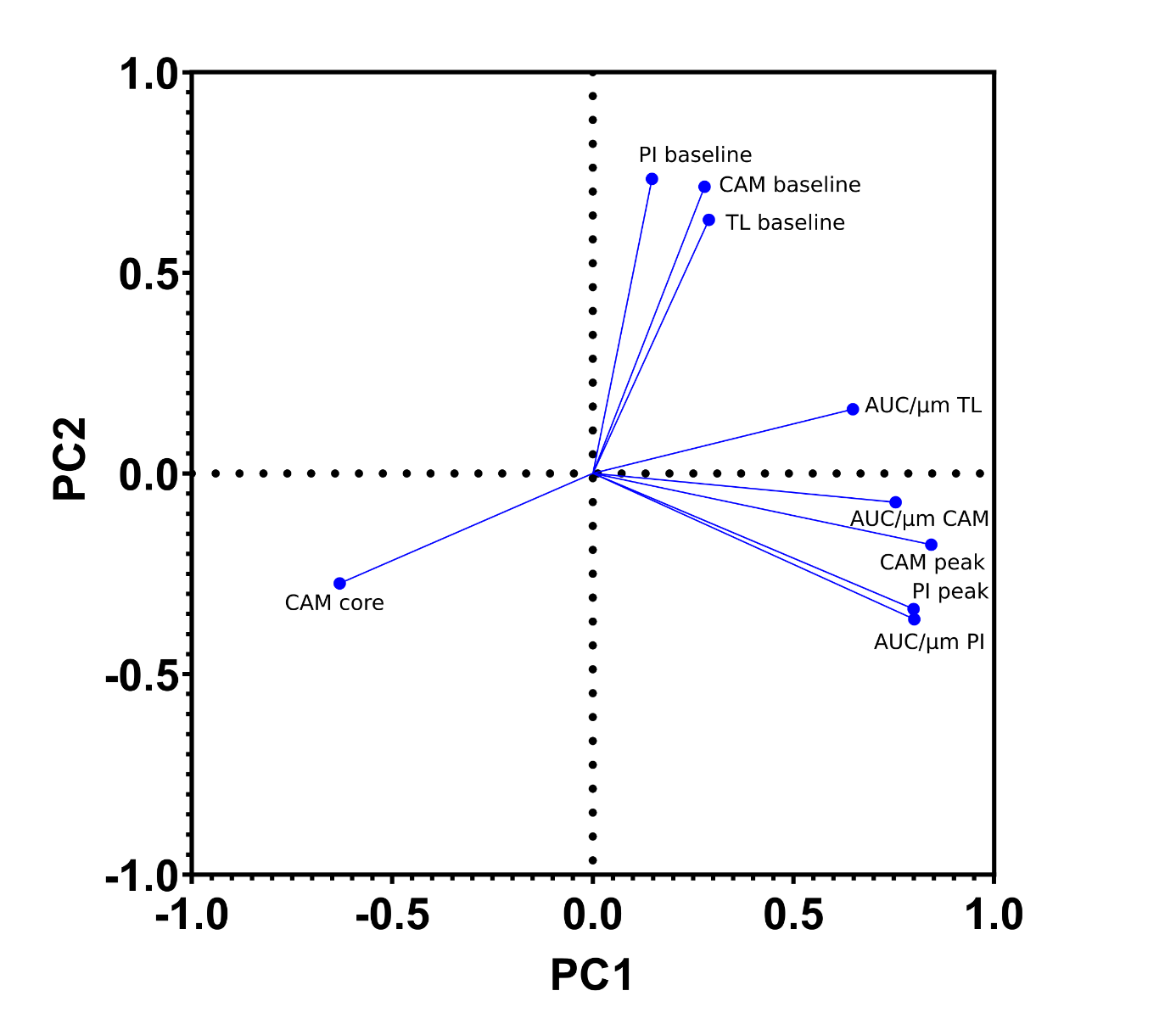

**Figure S4**. Loadings from principal component analysis of viability metrics. This shows the relative weight of each variable contributing to the principal component score. AUC/micron measurements are large contributors to positive PC1 scores, while CAM core contributes most in the opposite direction for PC1. The baseline values are collectively responsible variable for positive PC2 values while PI peak, AUC/micron, and CAM core weigh in the opposite direction. Abbreviations: calcein AM, CAM; propidium iodide, PI; transmitted light, TL; principal component, PC.

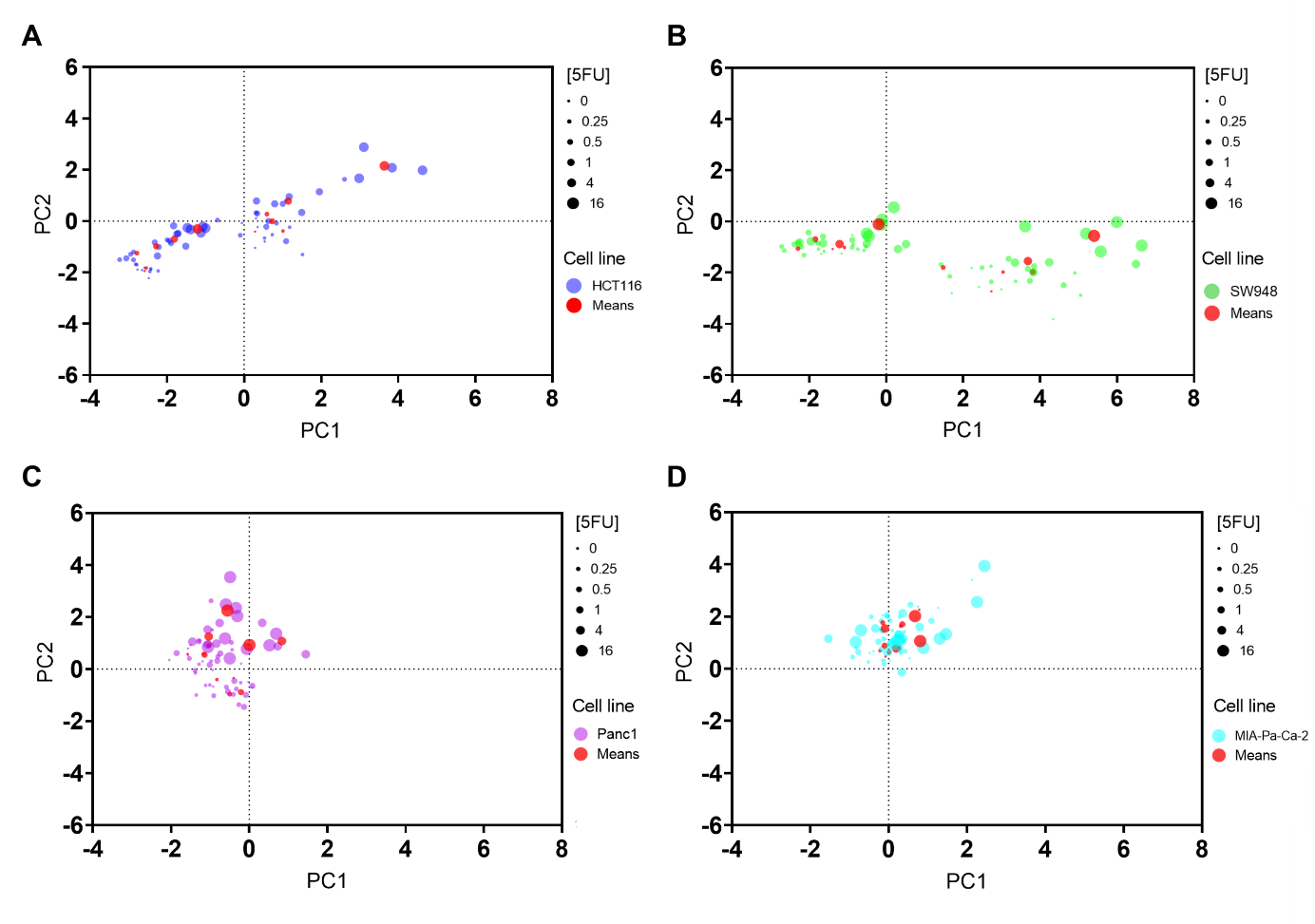

**Figure S5**. Principal component (PC) scores by cell line. (A) HCT116, (B) SW948, (C) Panc1, (D) MIA-Pa-Ca-2. Red dots are mean scores of 5-FU treatment groups.

**Table S1.** OCR/ECAR ratio of each cell line in 2D and 3D culture, from Tidwell *et al^28^*. In short, cultures were analyzed using a Seahorse XF96 instrument to probe metabolic phenotype. Values are from basal readings in the presence of DMEM culture media supplemented with 5 mM glucose, 2 mM L-glutamine.

| **Cell line** | **2D** | **3D** |
| --- | --- | --- |
| HCT116 | 1.196 | 0.871 |
| SW948 | 1.511 | 1.451 |
| Panc1 | 1.508 | 1.030 |
| MIA-Pa-Ca-2 | 1.673 | 1.166 |
