## Supplementary material for "Comparing *in vitro* cytotoxic drug sensitivity in colon and pancreatic cancer using 2D and 3D cell models: contrasting viability and growth inhibition in clinically relevant dose and repeated drug cycles": StackProfileData3

// StackProfileData 3.3
// This ImageJ macro gets the profile of all slices in a stack
// and writes the data to the Results table, one column per slice.
//
// Version 1.0, 24-Sep-2010 Michael Schmid
// Version 3.2, 07-Apr-2021 Tia Tidwell 
// specific to exported data with stack for each channel
//  --load 3 channel files first-- run for each set of stacks
// data output matches same structure as 2.0 macro
// Version 3.3, 09-Apr-2021: added set scale dialog


macro "Stack Profile Data" {

Odir = getDirectory("Choose Output Directory");
	
Dialog.create("Image position# to select windows for channel merging")
	Dialog.addString("Position #:", 01);
	Dialog.show();
	pos = Dialog.getString();


Dialog.create("Type and Set Scale")
	Dialog.addNumber("Number of pixels/um", 0.44);
	Dialog.show();
	n = Dialog.getNumber();

	run("Set Scale...", "distance=n known=1 pixel=1 unit=um global");

	selectWindow("Position0" + pos + "_RAW_ch00.tif");
	run("Stack to Images");
	selectWindow("Position0" + pos + "_RAW_ch01.tif");
	run("Stack to Images");
	selectWindow("Position0" + pos + "_RAW_ch02.tif");
	run("Stack to Images");

	run("Images to Stack", "name=Stack title=[] use");

	title = "WaitForUserSeelection";
 	 msg = "Make a line or rectangle selection, then click \"OK\".";
  	waitForUser(title, msg);

     if (!(selectionType()==0 || selectionType==5 || selectionType==6))
       exit("Line or Rectangle Selection Required");
     
     run("Plot Profile");
     Plot.getValues(x, y);
     run("Clear Results");
     for (i=0; i<x.length; i++)
         setResult("x", i, x[i]);
     close();

     n = nSlices;
     for (slice=1; slice<=n; slice++) {
         showProgress(slice, n);
         setSlice(slice);
         profile = getProfile();
         sliceLabel = toString(slice);
         sliceData = split(getMetadata("Label"),"\n");
         if (sliceData.length>0) {
             line0 = sliceData[0];
             if (lengthOf(sliceLabel) > 0)
                 sliceLabel = sliceLabel+ " ("+ line0 + ")";
         }
         for (i=0; i<profile.length; i++)
             setResult(sliceLabel, i, profile[i]);
     }
     setBatchMode(false);
     updateResults;
     selectWindow("Results");
		saveAs("Results", ""+ Odir + "Pos" + pos + "_results.txt");
	 run("Close All");
}
