## Supplementary material for "Comparing *in vitro* cytotoxic drug sensitivity in colon and pancreatic cancer using 2D and 3D cell models: contrasting viability and growth inhibition in clinically relevant dose and repeated drug cycles": SpheroidArea

// -------------------------------------------------------------------
// Written by: Tia Tidwell
// Date: Mar-2021
// Location: University of Stavanger, Stavanger, Norway
// New Location: ClexBio, Oslo, Norway
// Contact:
// -------------------------------------------------------------------

/*
This macro automatically detects and measures spheroids in a folder of images (optimized for brightfield).
The user to specify input folder (containing only images) and output folder for saving results. 
Results in the output folder include images displaying outlines of measured 
particles and a text document with particle (spheroid) detection information.  
Object measurements such as minimum and maximum particle size, area, etc. are 
determined manually by the user and entered at the prompts provided by the macro.
Specify these by using Analyze > Set Measurements command before running the macro to specify the measurements 
that are recorded for each object. 
*/ 


macro Spheroid_Area {

//Displays prompt for setting scale.  	
Dialog.create("Type and Set Scale")
	Dialog.addNumber("Number of pixels/unit", 440);
	Dialog.addChoice("Unit:", newArray("mm", "pixel", "um"));
	Dialog.show();
	n = Dialog.getNumber();
	u = Dialog.getChoice();
	
		
//Displays prompt for entering predetermined parameter values
Dialog.create("Parameters")
	Dialog.addChoice("Remove Outliers color:", newArray("Dark", "Bright"));
	Dialog.addNumber("Remove Outliers radius-1:", 2);
	Dialog.addNumber("Remove Outliers radius-2:", 5);
	Dialog.addNumber("Remove Outliers radius-3:", 10);
	Dialog.addNumber("Remove Outliers radius-4:", 10);
	Dialog.addCheckbox("Watershed", false);
	Dialog.addNumber("Analyze Particles - Min size:", 0.02);
	Dialog.addNumber("Analyze Particles - Max size:", 4);
	Dialog.addNumber("Threshold - lower limit:", 0);
	Dialog.addNumber("Threshold - upper limit:", 50);
	Dialog.show();

//Assigns the entered values to variables
	col = Dialog.getChoice();
	or1 = Dialog.getNumber();
	or2 = Dialog.getNumber();
	or3 = Dialog.getNumber();
	or4 = Dialog.getNumber();
	wtr = Dialog.getCheckbox();
	mncr = Dialog.getNumber();
	mxcr = Dialog.getNumber();
	tlo = Dialog.getNumber();
	tup = Dialog.getNumber();

//Displays Prompt for selection of Input & Output Directory
Idir = getDirectory("Choose Input Directory ");	
Odir = getDirectory("Choose Output Directory");


list = getFileList(Idir);
if (getVersion>="1.40e")
        setOption("display labels", true);
setBatchMode(true);
for (i=0; i<list.length; i++) {
        showProgress(i, list.length);
	processFile(Idir, Odir, list[i]);
    		}

function processFile(Idir, Odir, filename)
	{
	open(Idir + filename);
	run("Set Scale...", "distance=n known=1 pixel=1 unit=u global");
	run("Set Measurements...", "area mean min center perimeter fit shape area_fraction display add redirect=None decimal=3");

	//Convert RGB to 8-bit 
	run("8-bit");

	//Remove background by thresholding
	setAutoThreshold("Minimum");
	setThreshold(tlo, tup);
	run("Convert to Mask");
	run("Close-");

	//Closes and Fills Holes. Remove outliers step is for denoising and eliminating 	
	//debris particles. Can set size 0 in prompt if step is unnecessary. Watershed command separates fused 	//cells/colonies
	run("Remove Outliers...", "radius=or1 threshold=50 which=col");
	run("Remove Outliers...", "radius=or2 threshold=50 which=col");
	run("Remove Outliers...", "radius=or3 threshold=50 which=col");
	run("Fill Holes");
	if (wtr==true){
		run("Watershed");
		}
	run("Remove Outliers...", "radius=or4 threshold=50 which=col");

	//Analyze particles to measure highlighted objects. Minimum and Maximum sizes,
	//and circularities can be chosen in the prompt.
	roiManager("Reset");
	roiManager("Show All with labels");
	roiManager("Show All");
	run("Analyze Particles...", "size=mncr-mxcr show=Outlines display exclude clear include add");
		
	//Sends outlines from processed binary image to the original image via the ROI manager. Saves the original image displaying outlines in the output directory. 

	if (roiManager("Count") > 0){
		open(Idir + filename);	
		run("From ROI Manager");
		roiManager("Show All with labels");
		roiManager("Show All");
		Opath = Odir + filename;
		saveAs("JPEG", Opath);
		selectWindow("Results");
		saveAs("Results", ""+Odir + filename + "Results.txt");
		close();
	}
	
	run("Close All");
	
	}

}
